# Characteristics of Tactile Orientation Perception: Oblique Effect, Active vs Passive Exploration, and Serial Dependence

**DOI:** 10.1101/2025.03.19.644099

**Authors:** Guandong Wang, David Alais

## Abstract

Orientation perception is one of the most fundamental aspects of somatosensory perception, similar to its visual counterpart. In the current study, we examined several features of tactile orientation perception and their interactions, aiming to better understand the underlying mechanisms. We found that active exploration exhibits better orientation acuity compared to passive exploration. We also observed an exploration-independent and paradigm-independent robust anisotropy (i.e., tactile oblique effect) in tactile orientation acuity where the proximal-distal axis demonstrates superior orientation acuity compared to oblique and medial-lateral orientations. Finally, we demonstrated that, similar to its visual counterpart, tactile orientation perception showed a systematic attractive serial dependence effect, in which current orientation perception is biased towards the previous trial. However, this attractive effect is driven by post-perceptual processing/response rather than the stimulus *per se*.

## Introduction

Orientation perception is one of the most fundamental aspects of the somatosensory system. Tasks such as object manipulation (Pruszynski & Johansson, 2014) and shape perception (S. Hsiao, 2008) heavily rely on the ability to resolve orientation through touch. Similar to visual orientation perception, orientation information is extracted very early in the somatosensory processing hierarchy, with selectivity to edge orientations being found in first-order peripheral tactile neurons innervating the glabrous skin (Pruszynski & Johansson, 2014), and orientation information is well represented in the population response of primary (S1) and secondary (S2) somatosensory cortex neurons (S. S. Hsiao et al., 2002). Considering the aforementioned similarities, it is not surprising to suggest that the two modalities might share the same orientation processing mechanism for efficiency. Indeed, over the years, evidence from both neuroimaging (Nikbakht et al., 2018; Sathian & Zangaladze, 2002; Sathian et al., 1997; Zangaladze et al., 1999; H. Zhang & Alais, 2020) and behavioural studies (Lunghi & Alais, 2013; van der Groen et al., 2013; Wang & Alais, 2024) has accumulated in support of this hypothesis. Therefore, it can be imagined that more visual phenomena could be replicated in the somatosensory modality if they indeed share common processing mechanisms. This could also provide further insight into orientation processing in somatosensation.

The oblique effect is a well-known anisotropy in visual perception, where performance in various tasks is superior along the cardinal directions compared to oblique orientations (Appelle, 1972; Essock, 1980; Heeley et al., 1997). In recent years, evidence has accumulated to reveal the mechanism of this visual orientation anisotropy: it appears that what led to the superior acuity at cardinal orientations is the greater amount of orientation-selective neurons tuned to these orientations and the tighter orientation tuning of these cardinally-tuned neurons (Li et al., 2003). Moreover, the similarity between the pattern of this orientation anisotropy and the orientation distribution in natural visual scenes seems to suggest that an environmental prior is embedded in cortical processing through this mechanism, which enables efficient encoding of orientation information at a very early perceptual stage (Girshick et al., 2011; Harrison et al., 2023).

Inspired by the anisotropy in visual processing and the similarity in orientation processing between vision and somatosensation, researchers began investigating whether there is evidence for an analogous “oblique effect” in touch. Not surprisingly, a similar anisotropy has been demonstrated in tactile orientation perception (Essock et al., 1997, 1992; Gentaz & Streri, 2004; Gibson & Craig, 2005; Lechelt, 1988, 1992). Neurophysiological studies also showed that the slowly adapting type 1 (SA1) neurons, thought to be the main processing system of orientation information (S. S. Hsiao et al., 2002), is mildly biased towards stimuli in the proximal-distal orientation (Khalsa et al., 1998). However, unlike the rather consistent findings in vision, varied results have been reported regarding the somatosensory orientation anisotropy, with some reporting effects similar to the visual oblique effect where proximal-distal and media-lateral are more accurate than oblique (Lechelt, 1988, 1992), while others report proximal-distal is superior to other orientations (Essock et al., 1997, 1992; Gentaz & Streri, 2004; Gibson & Craig, 2005). Additionally, some studies reported no anisotropy in tactile orientation acuity (Craig, 1999). These mixed results raised an interesting question: Is this tactile oblique effect purely a result of the distribution of orientation-selection neurons, similar to the oblique effect in vision, or is it modulated by other factors that are unique to touch and which may vary between different tasks? One important difference between touch and vision that might mediate the observed anisotropy is the active nature of somatosensory perception.

Researchers over the years have spent considerable effort investigating how active movement affects tactile perception, revealing multiple differences between passive and active tactile perception. It has been shown that cutaneous inputs can be attenuated during voluntary movement of body parts, a phenomenon known as ‘tactile gating’. (Casado-Palacios et al., 2023; Chapman, 1994; Colino et al., 2014; Schmidt et al., 1990) By this principle active touch would be predicted to show inferior performance compared to passive exploration, however, most research comparing the two tasks showed little difference between the exploration methods (Olczak et al., 2018; Schwartz et al., 1975; Vega-Bermudez et al., 1991) or even active being superior to passive exploration (Heller, 1984; Smith et al., 2009). These contradictory results suggest that additional facilitatory mechanisms must be in play to counteract the effect of tactile gating. Complementing this, compared to passive touch, active exploration of a surface was found to produce different patterns of activation in the primary somatosensory cortex (S1) (Simões-Franklin et al., 2010), which shows that active exploration is not merely passive touch combined with additional motor movement. Instead, there appears to be a critical difference in cortical processes that differentiate these two modes of exploration, potentially engaging alternative sensory processing mechanisms and incorporating additional cues, which is likely to be developed as a result of our perceptual system adapting to the active nature of tactile exploration.

One possible origin of this difference is the additional cues involved during active touch due to the motor movement. Studies have found evidence that spatial (position) and intensity (force) cues are integrated during the active exploration of surface textures, combining in a manner consistent with maximum likelihood estimation (MLE) (Drewing & Ernst, 2006). The active movement of the finger introduces a changing temporal profile of receptor activation, which was found to improve performance on tactile spatial tasks (Cascio & Sathian, 2001; Gamzu & Ahissar, 2001).

Given the rather complex differences in processing mechanisms involved in active tactile perception, it is possible that the observed oblique effect could result from a combination of anisotropies originating at multiple levels of the tactile processing hierarchy. Hence the different processes in active and passive tactile exploration might lead to different patterns of the oblique effect. Hence, in Experiment 1, we implemented a factorial design to examine the impact of active versus passive touch and its potential interaction with the oblique effect. We hypothesised that the oblique effect may differ between active and passive exploration, and active exploration might reduce the potential benefit of the proximal-distal direction with the involvement of potential tactile gating and additional temporal information.

Another feature of visual perception that has been widely investigated in recent years is the serial dependence effect, in which the current percept is systematically biased towards the previous percept (for review, see Pascucci et al. (2023), Cicchini et al. (2024) and Manassi et al. (2023)). This effect was first demonstrated on orientation perception (Fischer & Whitney, 2014), and was then observed on a variety of visual tasks, from low-level features like numerosity (Fornaciai & Park, 2018) and colour (Barbosa & Compte, 2020), to high-level features such as face (Liberman et al., 2014), facial attractiveness (Xia et al., 2016) and aesthetic judgement (Kim et al., 2019). This attraction to perceptual history is thought to be developed by the brain as a means to facilitate stable perception against the constantly changing visual input Cicchini et al., 2018.

However, despite extensive investigations, a debate continues regarding the origin of this effect. Evidence from various studies supports the low-level perceptual (rather than cognitive or decision) nature of this bias, as the attractive bias is observed without previous decision (Manassi & Whitney, 2022; Murai & Whitney, 2021) or response(Fornaciai & Park, 2018). However, there are also contradicting findings that suggest that serial dependence requires conscious awareness of the previous stimulus (Kim et al., 2020), and gains in strength in post-perceptual processing such as when maintained in working memory for a later decision (Bliss et al., 2017). As the exact mechanism of the serial dependence effect is still unclear, more investigation is needed to clarify this puzzle.

Although a few attempts have been made to examine serial dependence in other modalities (Fornaciai & Park, 2019; Motala et al., 2020; Van der Burg et al., 2024), it has not been extensively investigated in touch. In our previous research, we were the first to demonstrate an attractive serial dependence effect in tactile orientation perception (Wang & Alais, 2024), as occurs in vision. However, as the paradigm was not optimized for serial dependence analysis, we undertook Experiments 2 and 3 to further clarify the serial dependence effect in touch. Given the similarity between vision and somatosensation in orientation perception, we proposed that a similar attractive serial dependence effect should be observed in tactile orientation perception. Consistent with this, Bae (2024) demonstrated that serial dependence in vision interacts with the oblique effect, which points to the two processes happening at a similar processing stage. Hence, we proposed to examine tactile serial dependence and whether it interacts with the oblique effect in Experiment 2. In Experiment 3, by separating the response from perception, we hope to investigate whether the observed serial dependence is indeed a low-level perceptual effect, or if it relies on the post-perceptual processing.

## Method

### Participants

A total of 36 naive participants were recruited for this study, 12 participants participated in each of the 3 experiments. All participants reported being right-hand dominant with no recent history of damage to the right index finger and no history of damage or diseases of the nervous system. This research was approved by the University of Sydney Human Research Ethics Committee (HREC 2021/048), and all methods were carried out in accordance with relevant guidelines and regulations. Participants were first and second-year psychology students from the University of Sydney, recruited through the Sydney University Psychology SONApsych research participation system, and were given course credit for their participation. Informed consent was obtained from all participants prior to the commencement of the experiment.

#### Apparatus and stimuli

The tactile stimuli used in all experiments were 3D-printed using an Ultimaker S5 3D printer. The circular stimulus disk had a diameter of 30 mm with the top surface being a triangular wave of 6 mm spatial period and a peak-to-trough amplitude of 2.5 mm, the disk was designed in FreeCAD and is shown in Figure. 1b. The stimulus disk was attached to a motor assembly that consisted of a stepper motor and a servo motor. Each step of the stepper motor corresponded to an angular rotation of 1.8^°^, and was used to control the orientation of the stimulus disk. The servo motor could elevate or lower the stimulus disk as commanded, which was used to control the presentation time of the stimulus. The motor assembly also contained a light sensor to calibrate the position of the stepper motor at the beginning of each test block. The motor assembly was controlled by MATLAB scripts via an Arduino UNO board and Adafruit Motor Shield V2.3. The aforementioned motor assembly was housed in a 3D-printed cuboid container, featuring a circular opening on the top surface with a diameter of 35 mm, through which the stimulus was raised for exploration by the participant. The entire box and motor device was fixed on a table in front of the participant. On the table beyond the box was a computer screen, which was used to deliver instructions to the participant during the experiment (See Figure. 1a).

**Figure 1.**
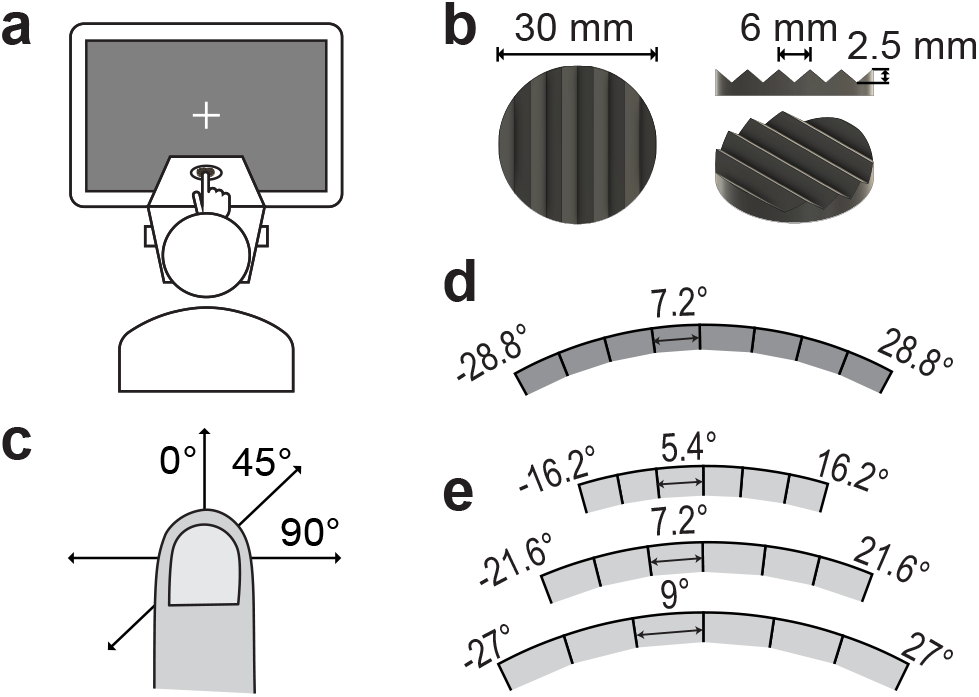
Experimental setup and the tactile stimulus used in the study. **a**. The participant was seated facing the tactile device, and a screen which displayed instructions, with their right index fingerpad positioned above the circular opening. The tactile grating disk was raised through the opening at a specified orientation and for a controlled duration. Participants were required to maintain an upright position and fixate on the fixation cross during the experiment to prevent body rotation. **b**. The tactile grating discs used in the experiment had a 30 mm diameter. They were 3D-printed with a triangular wave of 6 mm spatial period and 2.5mm peak-to-trough distance. **c**. Sets of stimulus orientations were centred around three orientations: proximal-distal: 0^°^, oblique: 45^°^and medial-lateral:90^°^. **d**. The nine stimulus orientations used in Experiment 1 ranged from −28.8^°^to 28.8^°^, with a 7.2^°^gap between each orientation level. These orientation levels are with respect to the centre orientation of each block. **e**. The three sets of stimulus orientations used in Experiments 2 and 3 were individually selected for each participant based on their performance in a preliminary test of orientation acuity. This approach aimed to address the significant variability in orientation acuity observed in Experiment 1, thereby ensuring better fitting psychometric functions for serial dependence analysis.

**Figure 2.**
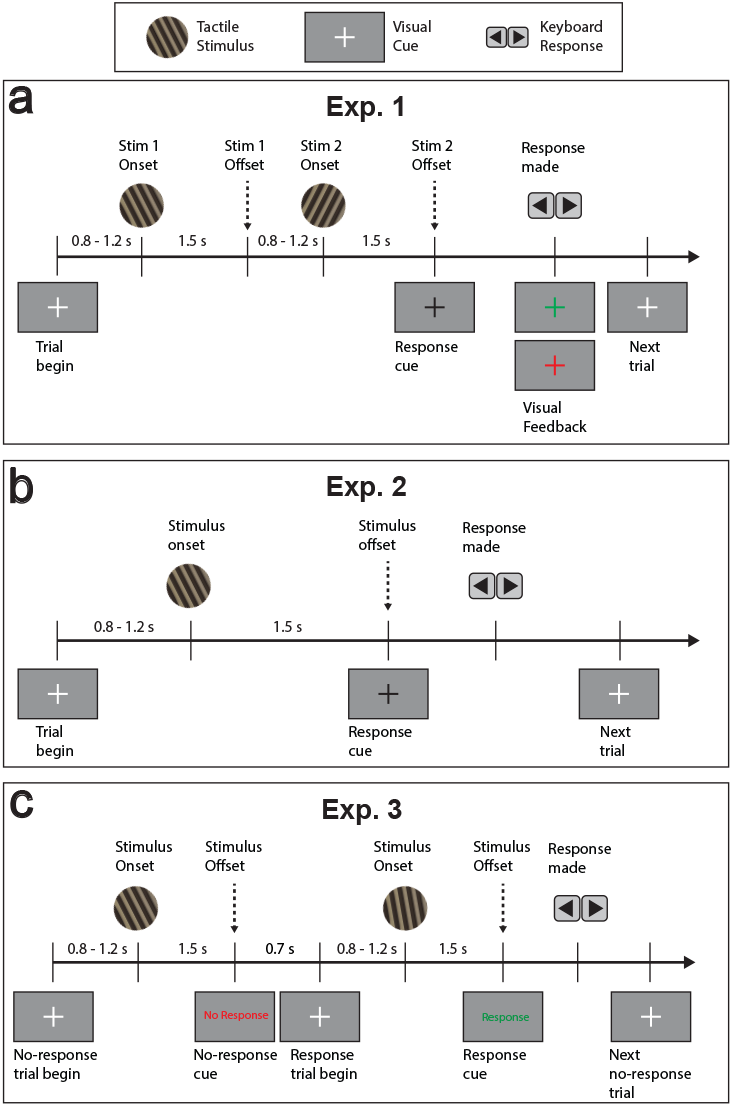
**a**. Experiment 1 Procedures: Each trial in the 2IFC task included two tactile stimulus presentations, each lasting 1.5 seconds. Following the presentations, participants responded by indicating whether stimulus 2 was CCW or CW relative to stimulus 1. Responses were made using arrow keys, and visual feedback was provided through a subsequent colour change in the fixation cross. **b**. Experiment 2 Procedures: Experiment 2 utilised a single-stimulus 2AFC task. After presenting a single stimulus, participants responded using the arrow key to indicate whether the stimulus is CCW or CW relative to a reference orientation. The reference orientations of the block (proximal-distal, oblique, or medial-lateral) were shown to participants prior to the commencement of each block. **c**. Experiment 3 Procedures: Experiment 3 employed the single-stimulus 2AFC task similar to Experiment 2, however, participants were cued to respond on only every second trial.

Prior to the commencement of each trial block, participants were instructed to sit in front of the screen with their heads resting on a chinrest. They were then asked to place their right hand on top of the container, with the right index finger over the circular opening and pointing straight forward towards the screen. At the beginning of each block, the grating stimulus disk was positioned underneath the top surface, which prevented the participant from touching it. During the experiment, the stimulus disk was rotated on each trial to the designated orientation by the stepper motor and then raised to the top surface through the circular opening by the servo motor so it could be touched (passively or actively, depending on condition) by the participant. It was then lowered back down after a controlled stimulus duration after each presentation.

### Design and procedure

#### Experiment 1: Tactile orientation acuity

The 12 participants involved in Experiment 1 attended two sessions on different days. In each session, they were instructed to perform the same orientation discrimination task using different methods (passive vs active), with the order of sessions counterbalanced to mitigate potential learning effects. For the passive condition, participants were instructed to passively feel the orientation of the stimulus disk without moving their index finger. In the active touch condition, participants were instructed to freely explore the grating disk by stroking their index finger over the grating to feel its orientation.

Each session of the experiment comprised six blocks. One of the three reference orientations (proximal-distal: 0^°^, diagonal: 45^°^, and medial-lateral: 90^°^, see Figure. 1c) was used for each block to assess the variations of orientation acuity at different axes. Each reference orientation was tested in two blocks. The sequences of the reference orientations were also counterbalanced between participants to minimize potential learning effects.

In both the active and passive conditions, participants were required to perform a simple two-interval-forced-choice (2IFC) orientation discrimination task. In each trial, the tactile grating disk will be presented to the participant twice, each lasting 1.5 seconds. The two stimuli consisted of the reference orientation of the block and a test stimulus, which was drawn from the nine test stimuli around the reference orientation, ranging from −28.8^°^to 28.8^°^with respect to the reference orientation of the block (see Figure. 1d). The order of the reference stimulus and test stimulus was pseudo-randomized to ensure an equal number of trials for both orders. The two presentations were separated by a randomized interval between 0.8 and 1.2 s as the stepper motor rotated the grating disk to the next orientation. The disk was rotated with a randomized component so that the interstimulus interval was not correlated with the orientation change so as not to cue participants about the orientation change.

After the two intervals of stimulus presentation, the fixation cross on the screen turned from white to black to cue the participant to respond using the left and right arrow keys, indicating whether the second presentation of the grating stimulus was rotated more counter-clockwise (left arrow key) or clockwise (right arrow key) in comparison to the first presentation. After their response, the fixation cross changed colour (correct response: green; incorrect response: red) to provide feedback and there was another 0.8 to 1.2 s random interval before the start of the next trial

Following the two presentations, the fixation cross on the screen changed from white to black, serving as a response cue. Participants indicated their response using the left and right arrow keys, signifying whether the second presentation of the grating stimulus was rotated more counter-clockwise (CCW: left arrow key) or clockwise (CW: right arrow key) compared to the first presentation. After the response, the fixation cross changed colour again (correct response: green, incorrect response: red) to provide visual feedback. After 300 ms, the fixation cross would turn white, indicating the start of the next trial.

#### Experiment 2: Tactile orientation serial dependence

As the two-interval 2AFC paradigm in Experiment 1 was not optimal for serial dependence analysis, Experiment 2 used a single-interval 2AFC task. The same three centre orientations used in Experiment 1 were used to replicate the oblique effect observed earlier and demonstrate the validity of the new paradigm as a measure of orientation acuity. However, in contrast to Experiment 1, only the passive touch method was employed.

Unlike Experiment 1, the range of the orientation used in Experiment 2 was different for each participant. At the beginning of each session, a preliminary test of approximately five minutes was conducted, which was designed to address the considerable variability in tactile orientation acuity observed in Experiment 1. The preliminary test employed the most extreme stimulus orientation from the three orientation sets to establish a preliminary estimation of participants’ relative tactile orientation acuity. One of the three stimulus ranges would be selected based on the participant’s performance in the preliminary test, to ensure a better psychometric function fitting for bias estimation (see Figure 1e). Moreover, each set of orientations only comprised seven levels to reduce the number of trials needed.

Prior to the commencement of the block, one of the three centre orientations was presented to the participant on the screen, serving as the reference orientation for that block. During each trial, participants were presented with a stimulus orientation for 1.5 seconds. Following this, they were visually cued by the colour change of the fixation cross and responded using the arrow keys, indicating whether the stimulus was more CCW or CW compared to the reference orientation of the block. No feedback was given in this experiment.

#### Experiment 3: Alternating response

To differentiate the serial effect between response and perception, we implemented an alternating response paradigm in Experiment 3. The procedure closely follows the design of Experiment 2, with only the proximal-distal reference orientation (0^°^) used in this experiment to further simplify the design. In this experiment, participants were cued to respond only every second trial. In the trials where no response was required, participants were instructed to solely focus on feeling the orientation of the stimulus without the need to make a response. Following the stimulus presentation, a “No Response” prompt appeared on the screen for 0.7 s, to maintain a similar time gap between trials as those with responses.

### Data analysis

#### Psychometric function fitting

Psychometric functions were individually fitted for each participant and each condition using the hierarchical Bayesian psychometric function fitting of the Palamedes Toolbox in MATLAB(Prins, 2023):

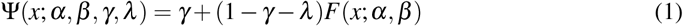

where *F*(*x*; *α, β*) is a cumulative Gaussian function with threshold *α* and slope *β*. The point of subjective equality (PSE) is obtained as the threshold (*α*, 50% CW response) of the fitted psychometric function, where non-zero PSE denotes the participants’ bias in orientation judgement. The slope (*β*) denotes the steepness of the psychometric function, which is used as a measure of orientation acuity. The parameters *γ* and *λ* represent the guess rate and the lapse rate, which denotes the difference between asymptotes of the psychometric function to 0% and 100%, respectively. The threshold (*α*) and slope (*β*) were set to vary freely across participants and conditions, while the guess rate (*γ*) was constrained to equal lapse rate (*λ*), and fixed across multiple conditions within a single participant. This was done to ensure that the guess rate and lapse rate can better capture the actual intentional lapse which should be roughly the same across different conditions as the conditions were broken into different blocks and counterbalanced within a session. Which should help better capture the potential differences in orientation bias and acuity among conditions through the two free parameters *α* and *β*.

#### Serial dependence analysis

The serial dependence effect was quantified as the difference between the point of subjective equality (PSE) between the different previous stimulus/response conditions, calculated as the equation below:

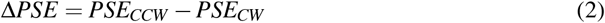

where a positive value indicates the current perception is attracted towards the previous stimulus/response and a negative value indicates the current perception is repulsed away from the previous stimulus/response.

#### N-back serial dependence effect

As the Shapiro-Wilk test indicated that the PSE differences deviated from normality, we used non-parametric permutation tests to analyse the n-back stimulus/response serial dependence effect. For each n-back stimulus/response condition, 10,000 permutations were created by random swap *PSE*_*CCW*_ and *PSE*_*CW*_ for each participant. The *p*-value of the permutation test is calculated as the proportion of permuted t-statistics that were greater than the observed t-statistic obtained from the original ΔPSEs (see Fig. 6c):

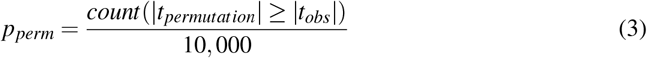

**Table 1.** Exp. 2 n-back statistics with respect to the previous stimulus.

| N-back | M | SD | $t_{obs}(11)$ | $p_{obs}$ | Cohen's $d_{obs}$ | $p_{Holm;perm}$ |
| --- | --- | --- | --- | --- | --- | --- |
| 1 | 6.505 | 5.383 | 4.186 | .002** | 1.208 | .0048** |
| 2 | 2.795 | 5.921 | 1.635 | .130 | 0.472 | .4210 |
| 3 | 0.507 | 4.192 | 0.419 | .683 | 0.121 | 1.0000 |
| 4 | 1.496 | 4.326 | 1.198 | .256 | 0.346 | 1.0000 |
| 5 | 0.599 | 2.190 | 0.947 | .364 | 0.273 | 1.0000 |
| 6 | -0.248 | 4.412 | -0.194 | .849 | -0.056 | .8606 |

*Note.* $p_{Holm;perm} > 1$ is set to 1.
\* $p < .05$ . \*\* $p < .01$ . \*\*\* $p < .001$ .

**Table 2.** Exp. 2 n-back statistics with respect to the previous response.

| N-back | M | SD | $t_{obs}(11)$ | $p_{obs}$ | Cohen's $d_{obs}$ | $p_{Holm;perm}$ |
| --- | --- | --- | --- | --- | --- | --- |
| 1 | 12.725 | 9.609 | 4.587 | < .001*** | 1.324 | .0032** |
| 2 | 7.048 | 7.539 | 3.238 | .008** | 0.935 | .0012** |
| 3 | 5.016 | 5.343 | 3.252 | .008** | 0.939 | .0035** |
| 4 | 4.006 | 4.016 | 3.456 | .005** | 0.998 | .0057** |
| 5 | 2.859 | 2.880 | 3.440 | .006** | 0.993 | .0058** |
| 6 | 1.323 | 3.378 | 1.356 | .202 | -0.392 | .2074 |

*Note.* $p_{Holm;perm} > 1$ is set to 1.
\* $p < .05$ . \*\* $p < .01$ . \*\*\* $p < .001$ .

The family-wise error rate was then controlled using the Holm-Bonferroni procedure:

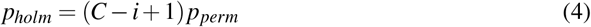

Where *C* represents the number of tests conducted for each family (in this case, *C* = 6 as the analysis was conducted up to the 6-back effect), and *i* represents the rank of the particular *p*-value in ascending order.

## Results

### Experiment 1: Tactile orientation acuity

#### Effect of touch method & reference orientation on orientation acuity

A two-way repeated-measures ANOVA was conducted to examine the effect of the touch method (passive, active) and reference orientation (vertical, oblique, horizontal) on the tactile orientation acuity (Figure. 3b). The tactile orientation acuity is quantified using the slope (*β*) of the fitted psychometric function, with a higher value (steeper slope) representing better orientation acuity. The sphericity assumption was checked for reference orientation (*χ*^2^(2) = 1.316, *p* = .518) and the interaction between touch method and reference (*χ*^2^(2) = 2.791, *p* = .248) using Mauchly’s test but neither was significant.

**Figure 3.**
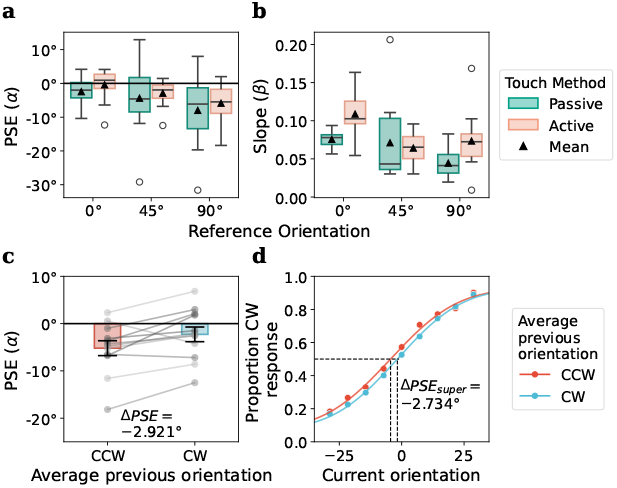
Experiment 1 results: **a**. The bias in orientation perception for all participants using the active vs passive touch method at the three different centre orientations (0^°^, 45^°^, and 90^°^) was quantified using the point of subjective equality (PSE) (threshold α from the fitted psychometric function, 50% CW response). A positive value represents a CCW bias, and a negative value represents a CW bias. No significant differences were found between active and passive touch (p = .228). Significant differences were found between different reference orientations (p = .031), with post hoc analysis showing the medial-lateral (0^°^) axis demonstrating CW biases compared to the proximal-distal axis (0^°^) (p_holm_ = .028) The central line represents the median, the box denotes the interquartile range (Q1–Q3), and whiskers extend to 1.5 times the IQR. Circles indicate outliers. And triangle indicates the mean. **b**. The tactile orientation acuity for all participants under each condition was quantified using the slope of the fitted psychometric function (β), where a higher value represents a steeper psychometric function (better orientation acuity). Active exploration showed significantly higher orientation acuity than passive touch (p = .006). When looking at the orientation acuity (slope of the psychometric function, β) is also significantly better near proximal-distal orientation (0^°^), compared to oblique (45^°^) (p_holm_ = .007) or medial-lateral (90^°^) (p_holm_ < .001). **c**. Serial dependence analysis: trials were divided based on the average orientation of the two stimuli in the previous trial, and the PSEs under the CCW and CW average previous stimulus orientations indicated a significant repulsive serial dependence effect (p = .012). Error bars represented standard errors. **d**. Psychometric functions for the supersubject data (all participant aggregated)reveal a shift in the PSE between the CCW and CW average previous stimulus orientation conditions. This shift indicates that the response in the current trial is repelled from the average orientation of the previous stimulus.

The passive touch condition (*M* = 0.064^°^, *SD* = 0.035) showed significantly lower tactile orientation acuity than the active touch condition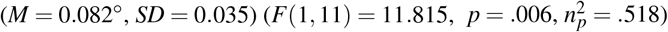. There is a significant main effect of reference orientation on tactile orientation acuity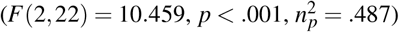. There is no significant interaction effect between the touch method and central orientation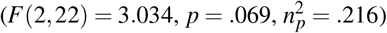. Post hoc tests on the main effect of centre orientation using Holm correction revealed that tactile orientation acuity was significantly higher for the stimulus around vertical orientation (*M* = .092, *SD* = 0.027) compared to the oblique orientation (*M* = .068, *SD* = 0.039, *t*(11) = 3.255, *p*_*holm*_ = .007, Cohen’s *d* = 0.781) as well as horizontal orientation (*M* = .059, *SD* = 0.033, *t*(11) = 4.410, *p*_*holm*_ *<* .001, Cohen’s *d* = 1.058). There is no significant difference between oblique and horizontal centre orientation (*t*(11) = 1.155, *p*_*holm*_ = .260, Cohen’s *d* = 0.277).

Another two-way repeated-measures ANOVA was carried out on the PSEs to examine the biases in tactile orientation perception under each condition. The sphericity assumption was checked for reference orientation (*χ*^2^(2) = 1.772, *p* = .412) and the interaction between touch method and reference (*χ*^2^(2) = 0.451, *p* = .798), but neither was significant.

The PSEs were found to differ significantly around different reference orientations 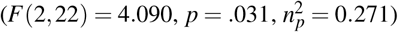. No significant differences in PSEs were found between passive ((*M* = *−*4.940^°^, *SD* = 9.280^°^) and active touch method 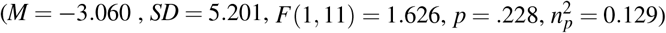. No significant interactions were found 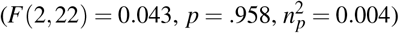.

Post hoc tests revealed that the orientation judgments near the horizontal orientation (*M* = *−*6.926, *SD* = 8.661) were significantly biased towards the CW direction compared to orientation judgments near the vertical orientation (*M* = *−*1.438, *SD* = 4.628, *t*(23) = 2.841, *p*_*holm*_ = .028, Cohen’s *d* = 0.744). No significant differences in PSEs were found between oblique (*M* = *−*3.637, *SD* = 7.915) and vertical (*t*(23) = 1.139, *p*_*holm*_ = .267, Cohen’s *d* = 0.298), or between oblique and horizontal (*t*(23) = 1.703, *p*_*holm*_ = .205, Cohen’s *d* = 0.446).

Although the 2AFC paradigm is not optimal for assessing the serial dependence effect, we attempted the serial dependence analysis by dividing the trials based on the average stimulus orientation of the previous 2IFC trial. The normality assumption was checked with the Shapiro-Wilk test, and no significant deviation from normality was found (*W* = .950, *p* = .636). A one-sample t-test on the ΔPSE revealed that trials with an average orientation of the previous trial CCW to the average of the current trial exhibited a significantly lower PSE compared to trials with an average orientation of the previous trial CW to the current trial (*M* = *−*2.921^°^, *SD* = 3.388, *t*(11) = *−*2.987, *p* = .012, Cohen’s *d* = −0.862), which indicates a significant repulsive serial dependence effect.

### Experiment 2: Tactile orientation serial dependence

To verify whether the oblique effect we observed in Experiment 1 was paradigm independent, two one-way repeated measures ANOVAs were performed to evaluate the effect of reference orientation on the PSEs (*α*) and slopes (*β*) with the single-stimulus test in Experiment 2. The sphericity assumption was checked for reference orientation for both analyses using Mauchly’s test and the assumptions were both met (PSE: *χ*^2^(2) = 2.795, *p* = .247; Slope: *χ*^2^(2) = 4.879, *p* = .087).

For the PSE analysis (Figure. 4a), unlike Experiment 1, no significant orientation bias was found at different reference orientations 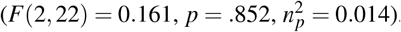.

**Figure 4.**
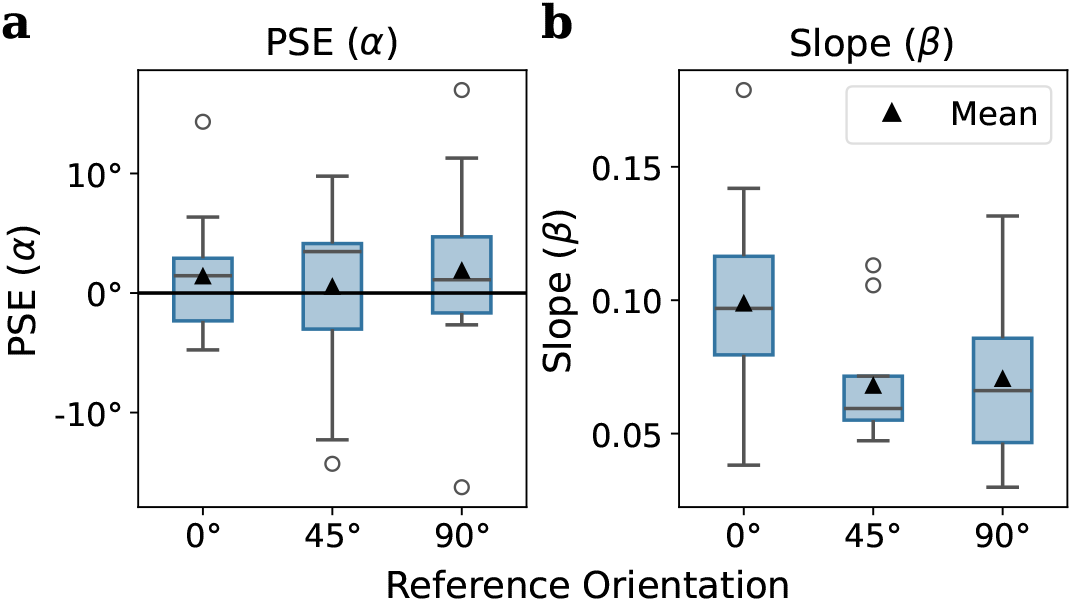
Experiemnt 2 results: **a**. PSEs (α) of the fitted psychometric functions at each reference orientation. Unlike Experiment 1, no significant differences were found between different reference orientations (p = .852). The central line represents the median, the box denotes the interquartile range (Q1–Q3), and the whiskers extend to 1.5 times the IQR. Circles indicate outliers. And triangle indicates the mean. **b**. Slopes (β) of the fitted psychometric functions at each reference orientation were used to assess the anisotropy in orientation acuity. Significant differences were found between different reference orientations (p = .021), with post hoc analyses indicating the proximal-distal axis (0^°^) exhibiting superior orientation acuity compared to the oblique orientation (p_holm_ = .035) as well as horizontal orientation (p_holm_ = .039).

When looking at the slope (Figure. 4b), orientation acuity differs significantly as a function of reference orientation 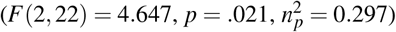. *Post hoc* analysis with Holm-Bonferroni family-wise correction found that orientation acuity at vertical orientation (*M* = 0.099, *SD* = 0.039) was significantly better than the oblique orientation (*M* = 0.068, *SD* = 0.021, *t*(11) = 2.747, *p*_*holm*_ = .035, Cohen’s *d* = 0.964) as well as horizontal orientation (*M* = 0.071, *SD* = 0.033, *t*(11) = 2.518, *p*_*holm*_ = .039, Cohen’s *d* = 0.883). No significant differences in orientation acuity were found between oblique orientation and horizontal orientation (*t*(11) = *−*0.229, *p*_*holm*_ = .821, Cohen’s *d* = *−*0.080).

To evaluate the serial dependence effect, a two-way repeated measures ANOVA was conducted to examine the effect of the previous stimulus orientation (CCW vsCW to the current stimulus) and reference orientation (vertical, oblique, horizontal) on the biases in tactile orientation perception (Figure. 5a). The orientation bias was quantified using the PSE (*α*) of the fitted psychometric function, where a positive value represents a leftward (CCW) bias in orientation, and a negative value represents a rightward (CW) bias. The sphericity assumption was tested for centre orientation (*χ*^2^(2) = 4.829, *p* = .089) and the interaction between previous stimulus orientation and centre orientation (*χ*^2^(2) = 1.199, *p* = .549) using Mauchly’s test, but neither was significant. Trials preceded by the CCW stimulus (*M* = 3.193, *SD* = 7.224) showed a significantly more leftward bias than the CW stimulus 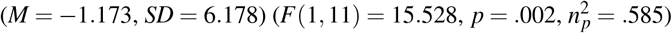, which indicates an attractive serial dependence effect. There was no significant main effect of centre orientation on the tactile orientation bias 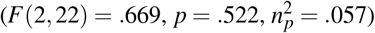 or interaction effect between the previous stimulus orientation and centre orientation 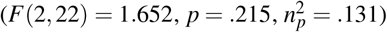.

**Figure 5.**
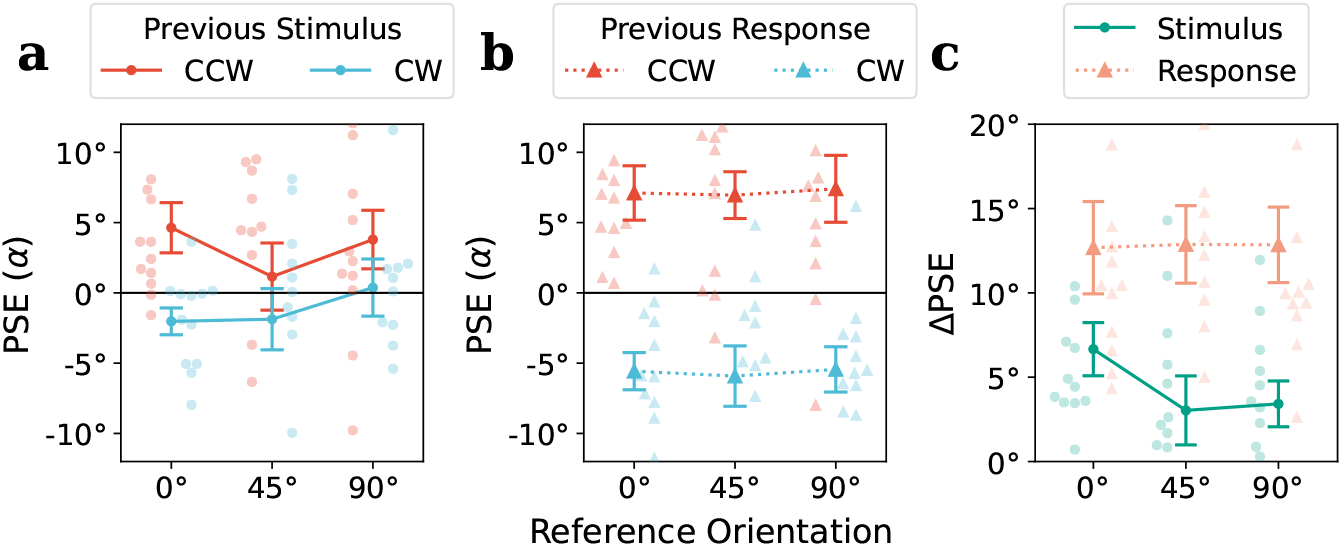
Experiment 2 serial dependence effect: **(a)** Significant attractive serial dependence effects were found with respect to the previous stimulus orientation (p = .002). No significant differences were found between different reference orientations, and no interaction was found between previous stimulus orientation and reference orientation. **(b)** Significant attractive serial dependence effects were found with respect to the previous response (p < .001). No significant differences were found between different reference orientations, and no interaction was found between previous stimulus orientation and reference orientation. **(c)** When comparing the stimulus-driven and response-driven serial dependence effect, the response-driven effect was significantly stronger (p < .001). No significant differences were found between reference orientations, and no significant interactions were found. Error bars represent standard errors.

**Figure 6.**
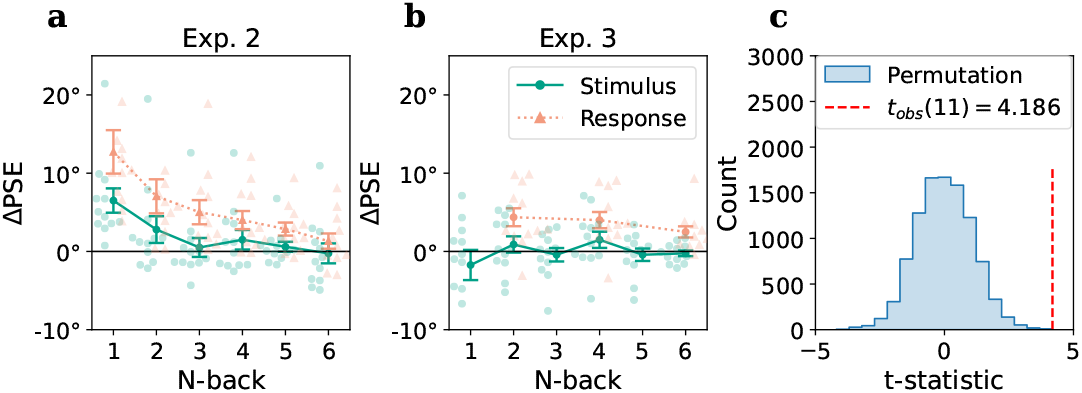
N-back serial dependence effect in Experiments 2 and 3: **a**. The mean ΔPSE between the trials with CCW stimulus and CW stimulus in the n-back trials (green line) and between CCW response and CW response in the n-back trials (orange line) in Experiment 2. Significant attractive serial dependence effects were observed for 1-back stimulus and 1-back to 5-back responses (see Table 1 and Table 2 for statistics). Error bars represent SEM. **b**. No significant serial dependence effects were observed with respect to the n-back stimulus. Significant attractive serial dependence effects were observed for 2-back, 4-back, and 6-back responses (see Table. 3 and Table. 4 for statistics). Error bars represent standard errors. **c**. An example of the non-parametric permutation test that was used to assess the n-back serial dependence effect. The example is for the 1-back stimulus in Experiment 2. The blue histogram shows the t-statistic distribution from the 10,000 permutations, the observed t-statistic is shown by the red dashed line. The p-value was calculated as the percentage of permuted t-statistics that exceed the observed statistics and were corrected using the Holm-Bonferroni procedure, p_Holm;perm_ = .0048.

A similar two-way repeated-measures ANOVA was conducted to examine the effect of the previous response and centre orientation on the biases in tactile orientation perception (Figure. 5b). The sphericity assumption was tested for centre orientation (*χ*^2^(2) = 2.246, *p* = .325) and the interaction between previous response and (*χ*^2^(2) = 1.055, *p* = .590) using Mauchly’s test, but neither was significant.

Trials preceded by the CCW response (*M* = 7.155, *SD* = 6.780) showed a significantly more leftward bias than the CW response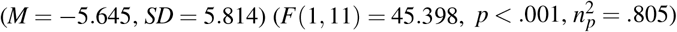, which also indicates an attractive serial dependence effect. There is no significant main effect of centre orientation on the tactile orientation bias 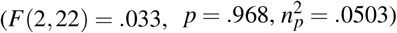 or interaction effect between the previous response and centre orientation 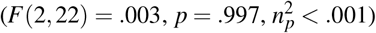.

To investigate the difference in attractive serial dependence effect between the previous stimulus and previous response (Figure. 5c), a two-way repeated-measures ANOVA was performed on the PSE differences (ΔPSE, see Eq. 2) of different type of previous influence (stimulus vs response) and centre orientations. The sphericity assumption was tested for centre orientation (*χ*^2^(2) = 0.261, *p* = .877) and the interaction between centre orientation and type of previous influence (*χ*^2^(2) = .873, *p* = .646) using Mauchly’s test, neither was significant.

The previous response (*M* = 12.800, *SD* = 8.189) showed a significantly more attractive serial effect compared to the previous stimulus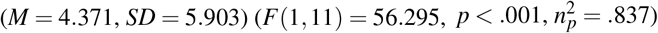. No significant differences were found for centre orientation 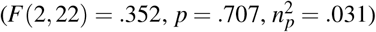. No significant interaction were observed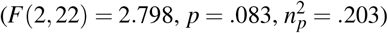.

### Experiment 3: Tactile orientation serial dependence with alternating responses

Shapiro-Wilk tests were carried out to assess the normality assumption and found the distribution of n-back ΔPSEs significantly deviated from normality for Experiment 2 (*W* = .779, .722, .724, .824, .778, .871, *p* = .005, .001, .001, .018, .005, .067). We therefore carried out multiple non-parametric permutation tests with family-wise Holm-Bonferroni correction to test the serial dependence effect with respect to n-back stimulus/responses for both Experiments 2 and 3.

To compare the results between Experiments 2 and 3, the analysis was only carried out on the blocks with a vertical reference orientation from Experiment 2. The attractive serial dependence effect with respect to the previous stimulus was only significant for the 1-back stimulus (see green line in Figue. 6a and Table. 1). With respect to the previous response, the attractive serial dependence effects were significant up to the 5-back response (see orange line in Figue 6a and Table 2).

In Experiment 3, no significant serial dependence effect was found with respect to the previous stimulus (see green line in Figue 6b and Table 3). Since the responses were only requested on every second trial, the attractive serial dependence effects can only be calculated for the 2, 4 and 6-back responses, and were found to be significant for all three (see orange line in Figue 6b and Table 4).

**Table 3.**
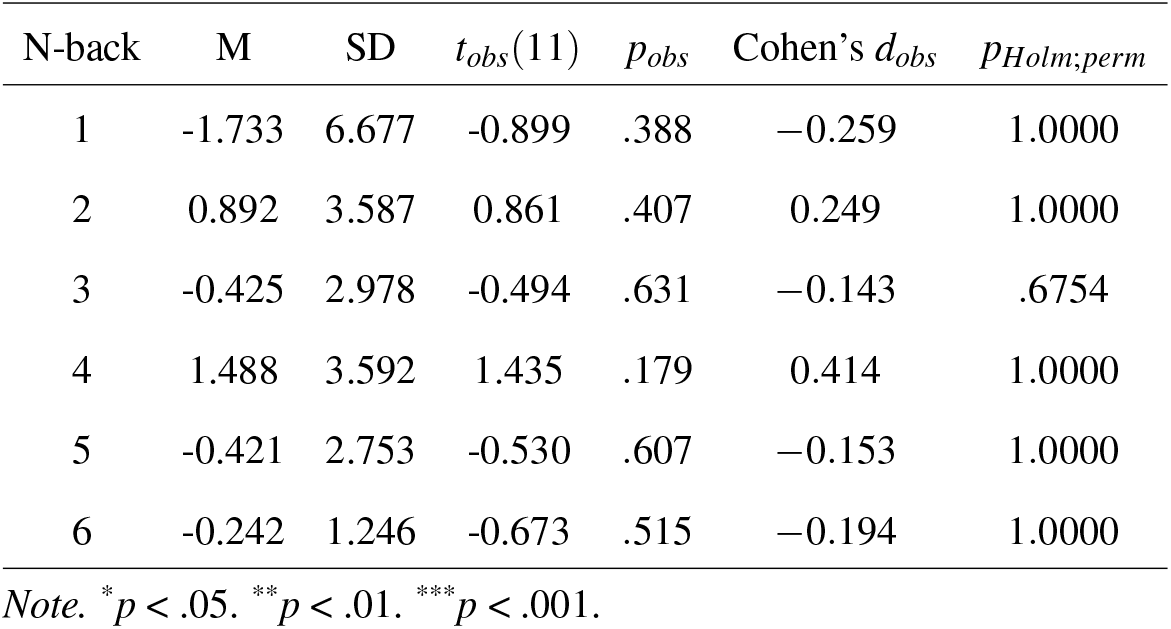
Exp. 3 n-back statistics with respect to the previous stimulus.

| N-back | M | SD | $t_{obs}(11)$ | $p_{obs}$ | Cohen's $d_{obs}$ | $p_{Holm;perm}$ |
| --- | --- | --- | --- | --- | --- | --- |
| 1 | -1.733 | 6.677 | -0.899 | .388 | -0.259 | 1.0000 |
| 2 | 0.892 | 3.587 | 0.861 | .407 | 0.249 | 1.0000 |
| 3 | -0.425 | 2.978 | -0.494 | .631 | -0.143 | .6754 |
| 4 | 1.488 | 3.592 | 1.435 | .179 | 0.414 | 1.0000 |
| 5 | -0.421 | 2.753 | -0.530 | .607 | -0.153 | 1.0000 |
| 6 | -0.242 | 1.246 | -0.673 | .515 | -0.194 | 1.0000 |

*Note.* \* $p < .05$ . \*\* $p < .01$ . \*\*\* $p < .001$ .**Table 4***Exp. 3 n-back statistics with respect to the previous response*

**Table 4.** Exp. 3 n-back statistics with respect to the previous response.

| N-back | M | SD | $t_{obs}(11)$ | $p_{obs}$ | Cohen's $d_{obs}$ | $p_{Holm;perm}$ |
| --- | --- | --- | --- | --- | --- | --- |
| 2 | 4.371 | 3.973 | 3.811 | .003** | 1.100 | .0046** |
| 4 | 4.003 | 3.596 | 3.856 | .003** | 1.113 | .0074** |
| 6 | 2.497 | 2.472 | 3.499 | .005** | 1.010 | .0033** |

*Note.* \* $p < .05$ . \*\* $p < .01$ . \*\*\* $p < .001$ .

We employed bootstrap methods to calculate means and confidence intervals, examining the impact of the correctness of the previous response and orientation on the serial dependence effect. In Experiment 2 (Figure. 7a), by dividing the previous trial based on its correctness, we can see that different patterns of serial dependence effect can be observed for correct and incorrect responses. For the trials where the previous trial’s response was correct (green), it shows an attractive serial dependence effect: when the previous stimulus was CCW, the bootstrapped PSE of the current trial was shifted towards the CW direction, meaning a CW stimulus was perceived as neutral, hence attracted towards the previous stimulus orientation, and *vice versa*. This attraction appears not to be affected by the previous stimulus orientation. On the contrary, the previous incorrect response showed a repulsive effect on the upcoming trial (red), in other words, the response of the current trial was repulsed away by the previous stimulus and attracted to the direction of the incorrect previous response. Although there seems to be increased variability between bootstrapped trials at the more extreme previous stimulus orientation, due to the decrease in a limited sample of incorrect trials with extreme stimuli, the bootstrapped mean and confidence interval still demonstrated a quite robust repulsion. In Experiment 3, the PSE was rather stable across all previous stimulus orientations when no responses were given (blue). These findings were in agreement with the previous analysis which shows that the attractive serial dependence effect was driven by the response.

**Figure 7.**
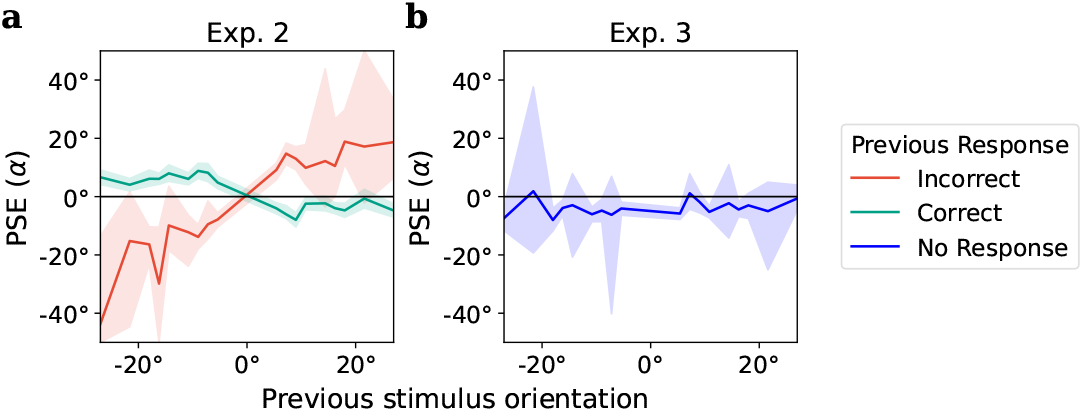
Bootstrapped estimates of the mean and 95% confidence intervals for the PSE across different previous stimulus orientations in Experiments 2 and 3: **(a)** Experiment 2: Trials were categorized based on the accuracy of the previous response. For each category, 10,000 simulations were performed to calculate the bootstrapped mean and confidence interval. Trials following a correct response (represented by a green line) demonstrated an attraction effect on the current trial, unaffected by the orientation of the previous stimulus. Conversely, trials following an incorrect response exhibited a repulsive effect on the subsequent trial (red), and this repulsion appeared to amplify with the orientation of the previous stimulus. **(c)** Experiment 3: the bootstrapped PSE seems to be rather stable across different previous stimulus orientations, where no responses were required (blue).

## Discussion

### Superior tactile orientation acuity at proximal-distal axis

Based on findings from the visual oblique effect (Girshick et al., 2011; Harrison et al., 2023), it is reasonable to hypothesize that the observed tactile oblique effect, where proximal-distal orientation shows significantly better tactile orientation acuity compared to oblique or medial-lateral orientation (Figure. 3b), may have a similar origin to its visual counterpart: This effect could originate from the orientation distribution of everyday tactile objects being embedded in the somatosensory system as a prior (analogous to natural scene statistics in vision (Hunt et al., 2009; Stansbury et al., 2013)), thereby facilitating efficient processing of orientation information. Our current findings seem to support this hypothesis. First, in Experiment 1, no interaction was found between the exploration method and the reference orientation, suggesting that the observed oblique effect is not influenced by the additional cues or tactile gating effect associated with active exploration. Therefore, it is more likely to be an anisotropy embedded in the actual cortical processing of orientation information that was shared between the different touch methods, rather than an artifact due to other factors such as movement patterns or finger shapes; Second, Experiment 2 showed when a single stimulus was present and compared with an internal reference system, the same pattern of oblique effect was observed (Figure. 4b), therefore further showing that the observed tactile oblique effect is not only touch-method independent but also paradigm-independent. The robustness of this tactile oblique effect demonstrated in our study, together with the neurophysiological evidence indicating that the orientation-encoding SA1 peripheral fibres’ responses were biased toward the proximal-distal orientation (Khalsa et al., 1998), seems to confirm the hypothesis of the orientation anisotropy being a built-in prior of the somatosensory system.

The evolutionary purpose of this embedded prior could be attributed to the efficiency of orientation encoding. Previous findings in both humans (S. S. Hsiao et al., 2002) and animals (Fitzgerald et al., 2006) suggest that the receptive fields of orientation-selective neurons may span multiple digits. This arrangement likely aids in orientation perception during object handling, as it is common for an edge or pattern on a tool or object to contact multiple fingers simultaneously. Given the structure of the human hand, an edge in contact with several digits is more likely to be oblique or medial-lateral relative to each finger pad rather than proximal-distal. Therefore, the superior orientation acuity at the proximal-distal orientation on a single finger pad could compensate for the lack of stimulation of neurons with cross-digit receptive fields, thereby creating a more efficient tactile orientation encoding schema. This hypothesis aligns well with the idea of anisotropy as an inherent prior within the perceptual system. Future studies could explore this possibility by testing orientation acuity across multiple digits.

Although the anisotropy of orientation acuity seems to fit with our hypothesis, upon investigation of the orientation biases, it was found that the PSEs also differed significantly around the reference orientations, with the horizontal orientation biased towards CW direction when compared with the vertical orientation (Figure. 3a). This finding was not anticipated given the 2IFC design and randomized sequence between reference and test stimuli in each trial. One possible explanation is that the observed bias could result from a recency effect, which describes the preference for more recent stimuli in a forced-choice paradigm (Mehrani & Peterson, 2015). In our Experiment 1, the observed clockwise (CW) bias reflects a tendency to favour the second stimulus over the first, particularly near horizontal orientations compared to vertical ones. This could be due to inferior orientation acuity near the horizontal axis, leading to increased uncertainty, where more trials fell below the just-noticeable difference. Consequently, participants were forced to guess, amplifying the recency effect in comparison to the vertical reference orientation. This would also explain why such a bias was not observed in Experiment 2. Despite a similar pattern of anisotropy in orientation acuity, each trial in Experiment 2 consisted of only a single stimulus presentation, thus preventing the systematic amplification of biases favouring either choice. This finding also indicated the importance of switching to the single stimulus design in Experiments 2 and 3, as the recency effect within each 2IFC trial would potentially mediate the investigation of the between-trial serial dependence effect.

### Active exploration shows better orientation acuity than passive

In Experiment 1, the active tactile perception showed a consistently better orientation acuity compared with passive tactile perception despite the reference axis (Figure. 3b). The advantage of active perception aligns with previous research (Heller, 1984; Smith et al., 2009), confirming that the disadvantage of potential tactile gating during active exploration can be compensated by the additional cue involved in active perception.

On top of this, numerous neurological studies indicate that active tactile perception is not merely an additive process to passive tactile perception. Evidence suggests differences in somatosensory cortical activation between the two exploration methods, likely mediated by top-down pathways (Krupa et al., 2004; Simões-Franklin et al., 2010).

In addition to the temporal cue, the active perception might also improve tactile orientation acuity via adaptation to the rapidly adapting (RA) afferents. Bensmaïa et al. (2006) found that the ratio between SA1 response and RA response correlated with the tactile spatial acuity, where RA response is likely to act as a form of noise during fine spatial perception. As the active movement during active perception involves the participants’ pad moving laterally across the grating surface, the mechanoreceptors are subject to vibratory stimulation, and since the RA fibre adapts quicker than the SA1 afferents (Leung et al., 2005), the reduced responsivity in RA afferents might help to amplify the ratio between SA1 and RA afferents, and hence improve the acuity in tactile spatial perception.

### Attractive serial dependence in touch is driven by the previous response

In Experiment 1, when separating the trials based on whether the average orientation of the previous 2IFC trial was CCW or CW to the average of the current trial, we noticed a significant shift in PSE that indicated a repulsive serial dependence effect (Figure. 3c,d). This repulsive serial effect seems to contradict the findings in visual orientation perception (Fischer & Whitney, 2014), as well as the attractive tactile serial dependence effect in our previous study (Wang & Alais, 2024). However, the two-interval 2AFC design while robust for investigating orientation acuity, is not optimal for serial dependence analysis due to its multiple stimulus presentations within one trial. Hence, in Experiment 2, we adopted a single stimulus paradigm to simplify the structure of each trial, and better reveal the influence of the previous trial. As previous research had indicated that serial dependence also can arise from a motor response and lead to individual differences in serial dependence (H. Zhang & Alais, 2020), the active exploration condition was removed from Experiment 2 to reduce the number of factors, and to keep the focus on serial dependence in tactile orientation processing.

Contrary to Experiment 1, an attractive serial dependence was found in Experiment 2. There is also no evidence of this attraction to the past being mediated by the reference orientation (Figure. 5a). These results seemed encouraging as they demonstrated an attraction to the recent past in tactile orientation perception similar to its visual counterpart. However further examination of the serial dependence effect with respect to the previous response showed an even larger attractive effect (Figure 5b, c). As the previous stimulus was unavoidably highly correlated with the previous response, it was unclear whether this attraction was driven by the stimulus *per se*, or by the post-perceptual processing/response.

To address this, in Experiment 3, we introduced an alternating response paradigm, where the participants were cued to respond on only every second trial. By doing so we were able to separate the serial dependence effect of stimulus and response. When looking at the n-back serial dependence effect, we found that there was no significant serial dependence effect with respect to the previous stimulus. However, the attraction to the previous response can be found to be significant up to the 6-back response. Compared to Experiment 2, where the stimulus-driven attractive serial dependence effect is significant for 1-back trials and the response-driven effect is significant for up to 5-back trials, these results further highlight a robust and long-lasting attractive serial dependence effect driven by response-related or post-perceptual processing.

To further examine the complicated interplay between stimulus and response in the previous trial, we plotted the bootstrapped mean and confidence interval across different previous stimulus orientations (Figure. 7). If the serial dependence effect were purely a perpetual effect, the PSE should not be affected by the correctness of the previous response. However, in Experiment 2, for the previous correct responses, attractive serial dependence was consistently observed across different levels of the previous stimulus orientations, with no obvious differences between different stimulus orientations. For the previously incorrect stimulus, a repulsive serial effect was found. This distinction between correct and incorrect previous responses indicated that the serial effect could not be a purely perceptual effect. The fact that attractive serial dependence (green line) seems rather consistent across the different previous stimulus orientations with a narrow confidence interval further supports the post-perceptual decision orientation of the attractive bias. However, for the previous incorrect response, the repulsive bias of PSE with respect to the previous stimulus suggested that the participant’s current response is biased towards the previous response, or in other words, their previous decision. Interestingly, this means and confidence intervals of the biases seem to also amplify as the previous stimulus is further away from the reference orientation. As the task in Experiment 2 was for the participant to determine whether the stimulus orientation was CCW or CW with respect to the reference orientation, the incorrect response with a more extreme stimulus seems to point more towards lapses where they missed the stimulus presentation. Hence the amplifying of the mean and confidence interval of the repulsive biases with respect to the previous stimulus could be a result of the participant’s tendency to unconsciously repeat the previous choice/motor responses when lacking information on the current trial.

Another possible explanation for the lack of serial dependence in Experiment 3 with respect to stimulus could also be the observed tactile orientation serial dependence effect originating from processing in the visual cortex. Various studies have found evidence for the involvement of the visual cortex during tactile orientation perception (Hidaka et al., 2022; Hu et al., 2021; Krystallidou & Thompson, 2016; Lunghi & Alais, 2013; Merabet et al., 2007; Sathian & Zangaladze, 2002; van der Groen et al., 2013; Zangaladze et al., 1999; M. Zhang et al., 2005). In our previous study, we also found that the tilt aftereffect (TAE) in touch could be transferred crossmodally to vision (Wang & Alais, 2024). Given the critical role of the visual cortex in tactile orientation perception, the observed serial dependence effect in Experiment 2 could also be manifest during orientation perception in the visual cortex. And as post-perceptual processing of orientation information is critical in activating the visual cortex (Zangaladze et al., 1999), the trials where responses were not cued might not involve the orientation processing in visual cortical areas, and hence did not induce an attractive serial dependence effect in the next trial. However, as the origin of serial dependence in visual processing and the connection between the visual and somatosensory cortices are still not fully understood, this remains a hypothesis.

## Conclusion

In this study, we conducted three experiments to investigate the intricate connection between the oblique effect, active exploration, and serial dependence in tactile orientation perception. By drawing analogies with visual orientation perception paradigms, we aim to better understand the tactile orientation processing mechanism and its similarity to visual orientation processing.

We demonstrated a robust tactile oblique effect that was independent of exploration methods (active vs passive) and paradigms (single-interval vs two-interval trials), with the proximal-distal axis showing superior orientation acuity compared to the oblique or medial-lateral axes. Additionally, we found that active exploration resulted in better orientation acuity across all reference axes. Finally, we demonstrated that the attractive serial dependence effect in touch is response-driven. These findings highlight the complexity of tactile orientation processing and its parallels with visual perception, offering new insights into the shared and distinct mechanisms underlying orientation processing.

